# Developmental Incongruity as a Dynamical Representation of Heterochrony

**DOI:** 10.1101/2020.07.31.231456

**Authors:** Bradly Alicea

## Abstract

The theory of heterochrony provides us with a generalized quantitative perspective on the dynamics of developmental trajectories. While useful, these linear developmental trajectories merely characterize changes in the speed and extent of growth in developmental time. One open problem in the literature involves how to characterize developmental trajectories for rare and incongruous modes of development. By combining nonlinear mathematical representations of development with models of gene expression networks (GRNs), the dynamics of growth given the plasticity and complexity of developmental timing are revealed. The approach presented here characterizes heterochrony as a dynamical system, while also proposing a computational motif in GRNs called triangular state machines (TSMs). TSMs enable local computation of phenotypic enhancement by producing nonlinear and potentially unexpected outputs. With a focus on developmental timing and a focus on sequential patterns of growth, formal techniques are developed to characterize delays and bifurcations in the developmental trajectory. More generally, growth is demonstrated using two conceptual models: a Galton board representing axial symmetry and a radial tree depicting differential growth. These techniques take into consideration the existence of multiple developmental genotypes operating in parallel, which ultimately characterize the exquisite phenotypic diversity observed in animal development.

## Introduction

Heterochrony [1,2] provides a framework for characterizing the nature of growth relative to adaptive changes in phenotype expressed in development and indicative of evolutionary history. More recent work on sequence heterochrony [3] reconsiders heterochrony as a series of developmental events such as tissue development or morphogenetic transformations positioned according to their relative occurrence. Taking a dynamical systems view of heterochrony [4] allows for the characterization of incongruous modes of growth and transformation during development [5] such as metamorphosis and compound growth. Incongruities exist in opposition to congruency over developmental time, which assumes that relative growth trajectories can be modeled with linear regression techniques. One example of a developmental incongruity is the metamorphic process, which results in a developmental trajectory that includes features such as delays and switching between stable developmental trajectories.

We can look to changes in neural organization that occur as a consequence of metamorphosis [6] to understand how heterochrony might operate in a system with variable dynamics in development. In organisms that undergo metamorphosis, structures that emerge in embryogenesis are broken down and rebuilt for the adult period. This results in two phases of development [7]: one driven by growth (construction of the embryo), and the other driven by reorganization and diapause (or suspended development). But in reality, what is referred to here as the suspension of development is a period of massive change in the phenotype driven by hormonal signaling [8,9]. Neural circuits follow this pattern: embryogenesis results in the formation of behaviorally irrelevant networks which are abruptly activated by changes during metamorphosis [10]. Metamorphosis also requires changes related to both neural cell bodies as well as changes in peripheral tissue (where neurons project their axonal processes) [11].

In terms of the rate of growth itself during metamorphosis, the number of cells can either stay the same or increase during this period of metamorphosis. In hemimetabolous insects, the neurons born during embryogenesis must be repurposed during metamorphic remodeling, as no additional cells are born post-embryonically [12]. By contrast, holometabolous insects exhibit two rounds of neurogenesis, which provides new cells for remodeling of the metamorphic phenotype [13]. In addition, metamorphosis is a mosaic process [14], affecting every cell differently. For example, cells can either retain the functional role acquired during embryogenesis, or act as polymorphic cells and acquire new functional roles [15]. This is characterized in this paper using a two-fold conceptual approach. The first is by using a Galton board (more popularly called a Plinko board) to simulate the accretion of a phenotype from a generator along an anatomical axis. This is done to demonstrate the accretion dynamics first observed in [16]. Behavior of the generator used in this exhibition is determined by our three parameters that describe dynamical heterochrony. This provides insights into how the mathematical modeling of heterochronic change maps to the real world. The second conceptual approach involves using a radial differentiation tree to generate tissues of different shapes and sizes using the parameters of dynamical heterochrony. Taken together, these examples will demonstrate a mathematical representation of biological growth and form.

### Biological relevance of Triangular State Machines

While the developmental phenotype can be characterized using a mathematical function, we must also represent a complex molecular state that underpins metamorphic change. We use a method called Triangular State Machines (TSMs) as a generative means to produce a phenotype. TSMs are embedded within a gene regulatory network (GRN) represented as a binary tree, and serve as a triangular motif incorporating a parent node, children nodes, and bidirectional connections between all three nodes. This representation enables the potential action of different genes in both *-cis* (acting within the reading frame, or in this case same level of binary GRN) and -trans (acting between loci, in this case across different levels of the binary GRN). As is generally the case for our GRN implementation, each node can be expressed at a rate between 0 and 1, which can be modified by activation in *-cis* and activation in -trans, and approximates a heterochronic function.

There are several governing rules for the operation of TSM structures. As triangular motifs, TSMs can be found anywhere in the structure of a binary GRN. However, only when TSMs exist in the lowest layer (tips of the tree) do they exhibit rules of decomposition and proportion. The rule of proportion states that the parent node (*A*) divides its value between its children *B* and *C*. Yet while *A* = *B* + *C, A* does not always equal *B*. It is also notable that when TSMs exist in upper layers of the binary GRN tree, child nodes can inherit full values (1.0) from the parent node. Upper layers can also exhibit lateral connections between children nodes to form a partial TSM. Every TSM can act in both *-cis* and -trans by means of a bidirectional lateral connection between child nodes *B* and *C*. The value of this lateral connection is an ascribed value proportional to the lowest valued child node. Finally, children nodes *B* and *C* can emit a phenotypic state by exhibiting a combinatorial rule. A child node can either act in *-trans* to express its full value, or act in *-cis* to express a fraction of its value weighted by the lateral connection between B and C.

As part of a binary GRN, TSMs collectively represent a phenotypic module that can emit phenotypic states along an anatomical gradient. When a TSM is embedded in the overlap between two GRNs, the output resembles the boundary between two morphogen gradients [17]. This can be demonstrated by the formation of Moire patterns, which are occasionally found in the morphogenesis of multicellular biological systems [18,19]. The combination of conditional expression of TSMs and overlapping GRNs are an important source of gene-gene interactions, in addition to connecting spatial heterogeneity to temporal dynamics. TSMs also serve as enhancers for upstream genes and their expression. A demonstration of TSMs and how they are embedded in GRN topologies is shown in Supplemental Movie 1.

The dynamical view of heterochrony defines these phenomena in two ways: dynamical multiphasic heterochrony and compound heterochrony. While heterochrony can be understood using the distilled mathematical representation, a genomic representation that emits phenotypic elements can also be used to generate a wide range of heterochronic patterns and new phases. Directed genetic regulatory networks (GRNs) determine the expression of heterochrony, and are represented in the form of binary trees. These binary GRNs can exchange information laterally and in some cases overlap with other such trees. Aside from functioning as a network of expressed genes that exhibits epistasis (interactions between genes), lateral connections and overlapping gene expression patterns form functional motifs composed of triangular state machines (TSMs). Through the output of GRNs composed of TSMs, one or more developmental trajectories are produced. The mode and control of these systems can be characterized using a dynamical systems approach [4], and can explain complexity in the developmental process. These outputs will be mapped to toy models of the phenotype in which growth is characterized by changes in size and/or shape.

### Acquisition of Developmental Sequences

One essential component of heterochrony is understanding the relationship between various developmental stages. According to Williamson [5], incongruous larval stages that resemble those of distantly-related taxa can be observed in several orders of marine invertebrate. From a sequence heterochrony point-of-view, the growth trajectory can be rearranged across phylogeny as developmental sequences are swapped and newly activated (Figure 1). This rearrangement of developmental sequences has consequences on the growth trajectory, from nonlinearities to discontinuities [20]. While we can gain insight from the notion of sequence heterochrony, current theoretical models do not explain developmental incongruities. The theoretical perspective described in this paper provides a clearer basis for how these incongruities might arise and be expressed in the course of development.

**Figure 1.**
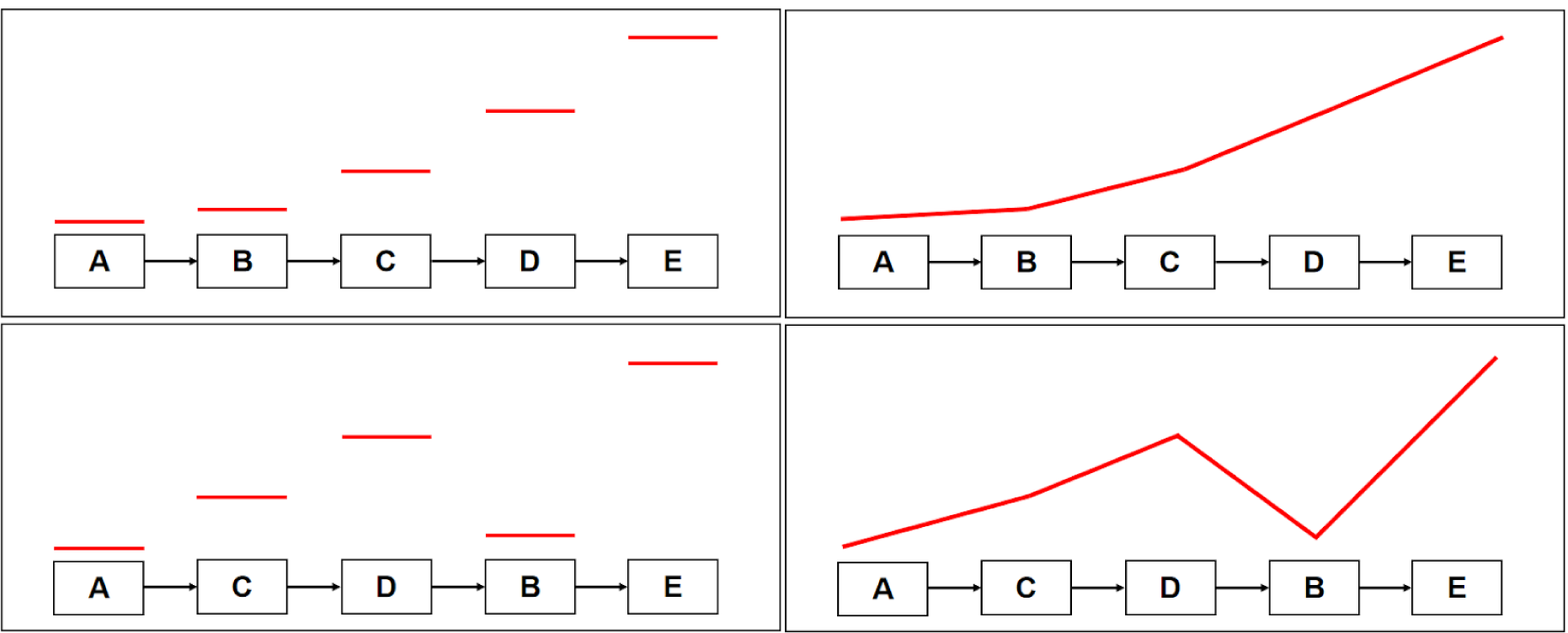
Diagram demonstrating the rearrangement of developmental sequences and its relationship to dynamical developmental trajectories. TOP: at left, initial sequence of developmental stages and their initial length (uniform); at right, their expression as a trajectory over time. BOTTOM: at left, rearranged sequence of developmental stages and their initial length (uniform); at right, their expression as a trajectory over time. Besides rearrangement, developmental sequences can also be contracted or dilated over time (not shown).

This model of heterochrony indirectly addresses whether multiple developmental stages accumulate through singular events or as a series of serial acquisitions. In the case of hybrids, combinations of genetic material from two independent species occur either in convergence zones [21,22] or as multiple independent interbreeding events resulting in reticulate evolution [23,24]. Furthermore, hybridization events can lead to both large-scale genomic rearrangements and changes in gene expression [25,26]. While horizontal gene transfer provides a more explicit mechanism [27-29], specific instances are no easier to empirically identify. Therefore, we should utilize models of both a genomic representation and a more mathematically sophisticated representation of heterochrony to better characterize this opaque phenomenon.

### Expression of Developmental Sequences

Heterochrony can be defined as changes in the rate of change in development for a specific trait, the resulting acceleration or deceleration being a result of tweaks made to the expression of genes in any given genetic regulatory network (GRN). In more complicated life-history trajectories such as metamorphosis, heterochrony does modify a linear sequence of events. Instead, heterochrony may tend to coordinate multiple developmental trajectories, such as the two types of overlapping body symmetry observed in the development of sea cucumbers [5, 20]. This overlap seems to be due to heterochronic changes in the expression of each type of symmetry relative to their ancestral expression: radial symmetry is accelerated and expressed in late larvae, while the disappearance of bilateral symmetry is decelerated so that it persists throughout adulthood. When considered as a rearrangement of developmental order, these might be called “start again” developmental sequences [20]. Ultimately, the model presented here might be helpful in evaluating such observations.

### Assumptions and Methods

One way to interpret the interactions of developmental sequences is by treating them as a dynamical system. Based on *Drosophila* embryogenesis, a number of studies [30-34] suggest that development can be characterized as a dynamical system rather than a series of discrete programs executed serially [4]. This view seems to conflict with conventional views of heterochrony, so a series of assumptions need to be made in order to bridge the gap between the language of heterochrony and developmental dynamics.

### Heterochrony

This first new concept is to extend the theory of heterochrony, including defining a mathematical formalism for dynamical multiphasic heterochrony and in particular the compound case. As an extension of the mathematical model of heterochrony proposed in [1], dynamical multiphasic heterochrony explains distinct phases of phenotypic complexity across developmental time, as well as the transition from one phase to the other. Compound heterochrony relies on three parameters: α, β, and τ. Consistent with the formalism in [1], α and β represent the initiation and termination of growth, respectively. The parameter τ is consistent with the notation of delay differential equations (DDEs), and represents the length of delay between two growth phases, each being a function defined by an α, β pair. The nature of these compound dynamics depends on the value τ, operationalized as the length of time between β_*t*_ and α _*t*+1_. When τ is positive, there are two distinct phases of growth in the organism. When τ is negative, the transition between phases is smoother. The specific trajectories of compound heterochrony also rely upon the parameter k, or the overall growth rate.

#### Dynamical Heterochrony Model

The growth law of [1] can be restated to represent differential growth over developmental time. Differential growth occurs under three conditions

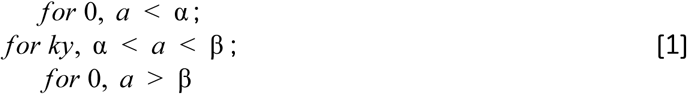

where *a* is one developmental window, and *y* grows according to Eq. 2 between onset age α and offset age β.

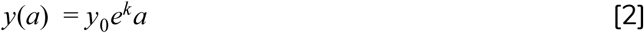

To bring together the simple model of heterochrony with a compound model, we can use Delay Differential Equations (DDEs). DDEs are characterized by time-delay systems in [35]. In their general form, a time-delay system is

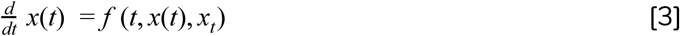

where *x*_*t*_ = [*x*(τ) : τ Δτ] represents the trajectory of a solution in the past. Reformulated as a delay differential equation (DDE) with a single delay, the equation can be structured as

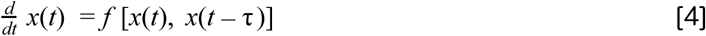

We can also characterize time delays more specifically with respect to heterochrony and developmental growth trajectories. In equation (1), α and β both exhibit systematic time delays (τ): α (τ), β (τ). Delay in the growth trajectory is characterized over the interval α, β, and is equivalent to Δ*k*. The total length of the delayed process is (β + τ) – (α + τ). Using this formulation, the rate of a delay process is *k*_0_, the delay rate is 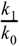, and the length of a delay process is β_*x*_ − α_*x*_. Additional mathematical and graphical details of these newly-identified forms of heterochrony are defined in Supplementary File 1, which provides details and mathematical formalisms, conceptual results, connections to the developmental constraints such as the critical period, and theoretical scenarios for nonlinear switches during developmental growth.

### Genetic Regulatory Network

A variant of the Cayley tree model described by Artyomov et.al [36] is used to approximate a suitable GRN. Each GRN (a single binary tree) represents a combination distinct phenotypic modules and expression of heterochrony genes [37] that exist to control the timing of developmental processes. Each level of the tree allows us to transform a single parent node into two children. This can either be one gene affected by another, or a gene affected by a regulatory element. In following a path between the root node to a branch tip, we encounter genes that have a direct influence of other genes (no intermediate regulatory elements), and genes that are influenced by regulatory elements (promoters, enhancers). There are also lateral pathways where one the effects of one genetic pathway can influence the effect of another (epistasis).

#### Binary GRN Model

The binary GRN used here can be defined as a binary tree *T* rooted at node *N* _0_ expressed at time *t*_0_. This node can be defined as a master control gene, which turns on children nodes (genes) *A* and *B* at time *t*_1_. Each of these children represent a gene under contingent upon the master gene. Subsequent layers of the tree represent genes that are dependent upon child nodes *A* and *B*, and act as promoter or enhancer elements (depending on their depth in the tree) that contribute to the phenotype. Growth parameters α, β, and τ can be used to control the timing of nodes (birth of children from the parent) and the execution of rules in the TSM. The degree of activity along all of these arcs must be pre-defined, while tips of the trees produce a continuous output that shape the level of growth at a particular point in time.

Generally, the master control motif exists at the top of the binary tree (at the root), promoters can exist in the intermediate layers (generally from *t*_2_ to *t*_*n*−2_), and enhancers exist among the bottom two layers of the tree (at the tips). In this non-overlapping example, one child node (*A*) is a gene of lesser effect, while the other child node (*B*) is a gene of greater effect. The expression of both genes are mediated by promoters. The gene of lesser effect (*g*_*0*_) has two promoters *p*: *p*_*00*_ and *p*_*01*_, while the gene of greater effect (*g*_*1*_) also has two promoters: *p*_*10*_ and *p*_*11*_. Each pair of promoters are expressed according to probabilities *p* and *1-p*, respectively. The relative probability of switching is context-dependent. Enhancers, present at *t*_*3*_ in this example or as the action of TSMs, mediate the expression of genes *0* and *1* as the children of a promoter node. In this example, each promoter would have two enhancer nodes, for a total of eight (8) enhancer nodes. Unlike promoters, enhancers act as analogue tuners, while each enhancer (child) produces a modified output of its parent (promoter).

To show how activation works and outputs are produced, a path can be traced through the tree from the root node to a specific branch tip. Paths between nodes have an activation value, which can either be in the form of binary switches (0 for off, 1 for on) or analogue dials with continuous values from 0 to 1 (for regulatory elements). An example of a switch is between a master control gene and one of its immediate children. When the value is 0, the effect of the master control gene is off for that regulatory relationship. These switches can also be used as a timing mechanism for the onset and offset of the growth function (e.g. α and β) [38].

Returning to the non-overlapping example, suppose the following: an open path between the master control gene and its children is 1.0, the path between that child and the promoter node is 0.3, and the path between the promoter and the enhancer node is 0.65. As a multiplicative process, this path produces an output of 0.195. The gene-gene pathway represents the co-activation of the two genes, while the regulatory elements allow for proportions of that co-activation to be expressed. The lateral connections allow for pathways to cross, combining inputs from lateral and horizontal pathways through multiplication. The branch tips of the tree can either be mapped to spatial locations in the phenotype at time *t* or summed across all branch tips to reveal the proportion of full activation produced at time *t*.

## Results

To further understand the dynamics of heterochronic processes, a distinction must be made between two alternate models for characterizing heterochrony in life-history: multiphasic and compound. Multiphasic heterochrony involves plastic responses to growth stimuli over developmental time. In an indirect manner, the multiphasic case relies upon sequence heterochrony (see Figure 1). In a similar fashion, compound heterochrony involves multiple developmental trajectories expressed within the same organism. Both of these types can also be characterized using a dynamical formulation of previous descriptions of heterochrony.

While metamorphosis is one example, another more common example are differential growth rates exhibited by different segments or modules of the same organism. This can be observed in taxonomic groups as diverse as arthropods [39] and humans [40]. The latter example is generally identified using allometric scaling relationships [41]. These types of heterochrony are demonstrated in Figures 2 and 3, respectively.

**Figure 2.**
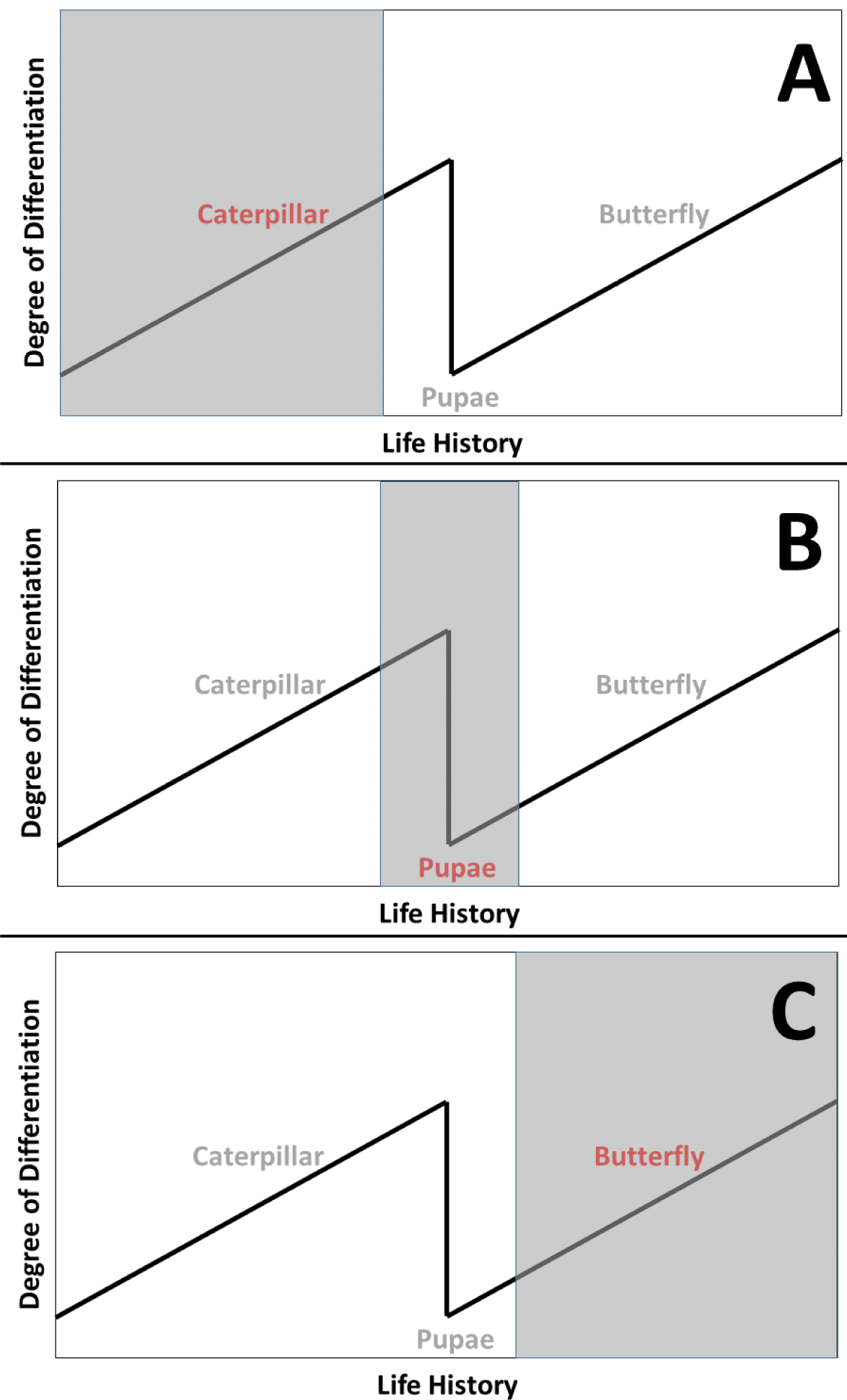
A step-by-step description of multiphasic heterochrony, in which the larval form (caterpillar) dedifferentiates into an intermediate pupae. The developmental trajectories of two heterochronic regimes expressed as two distinct parts of life history. A) the caterpillar heterochronic trajectory governs early in life-history, B) an intermediate diapause stage occurs where growth stops and selected portions of the phenotype are obliterated, C) the butterfly heterochronic trajectory governs later stages of life history.

**Figure 3.**
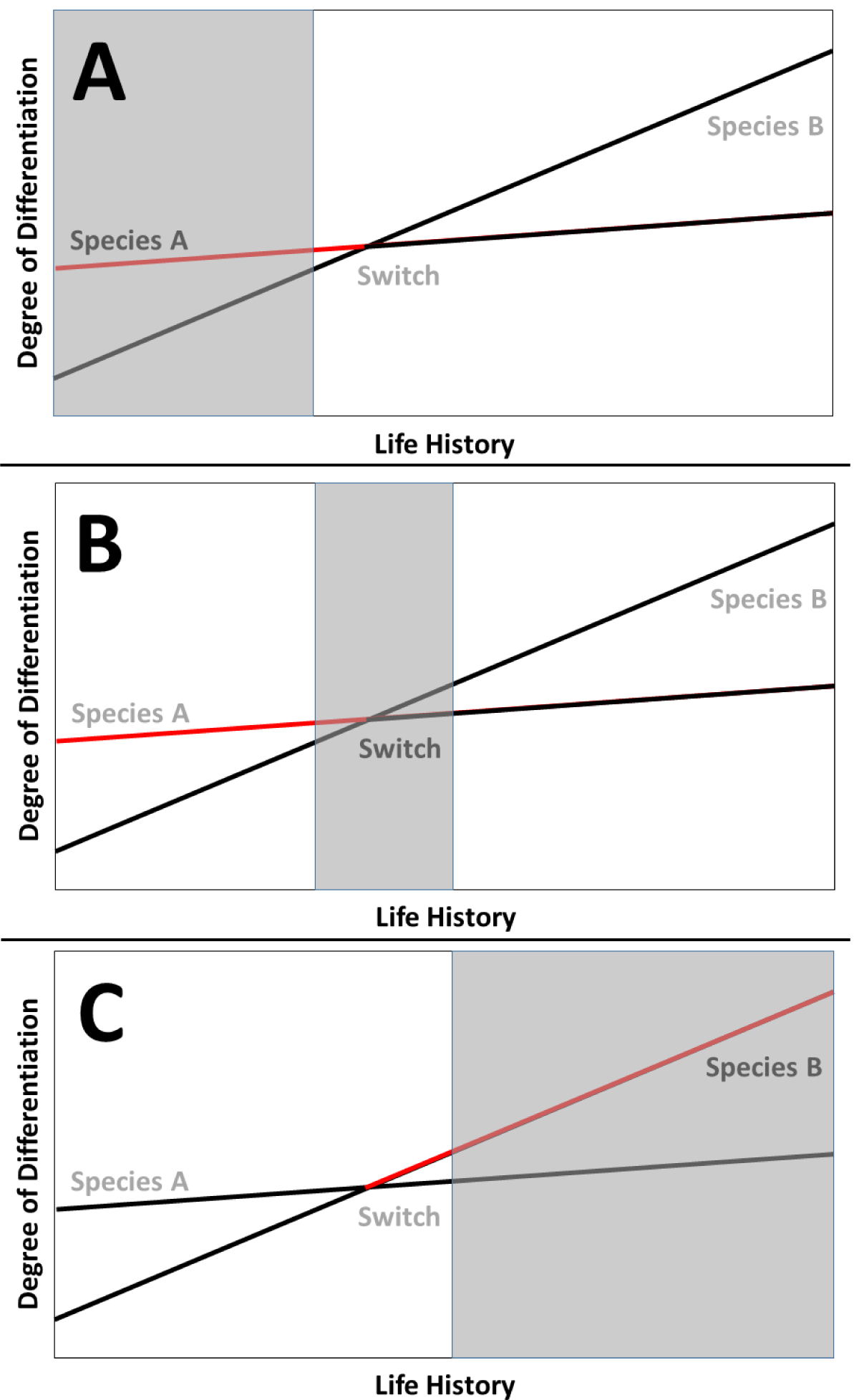
A step-by-step description of compound heterochrony, in which developmental trajectories from two different species overlap, resulting in a switch from one heterochronic regime to the other over the course of life history. A) the heterochronic trajectory from Species A is active early in life-history, B) a switch between heterochronic trajectories is activated, C) the heterochronic trajectory from species B is active later in life-history.

Figure 4 demonstrates a compound heterochronic process with a positive value for τ. This example confirms that compound trajectories also rely on different values for *k*, in addition to the difference in individual α and β points along each part of the greater developmental trajectory. In this case, Δβ is much greater than Δα, which suggests exponential growth in the later phases of development. As an α, β pair represents a single binary representation of phenotypic modules and heterochrony genes, τ is also related to the degree of overlap in adjoining binary GRNs and thus the size of a TSM.

**Figure 4.**
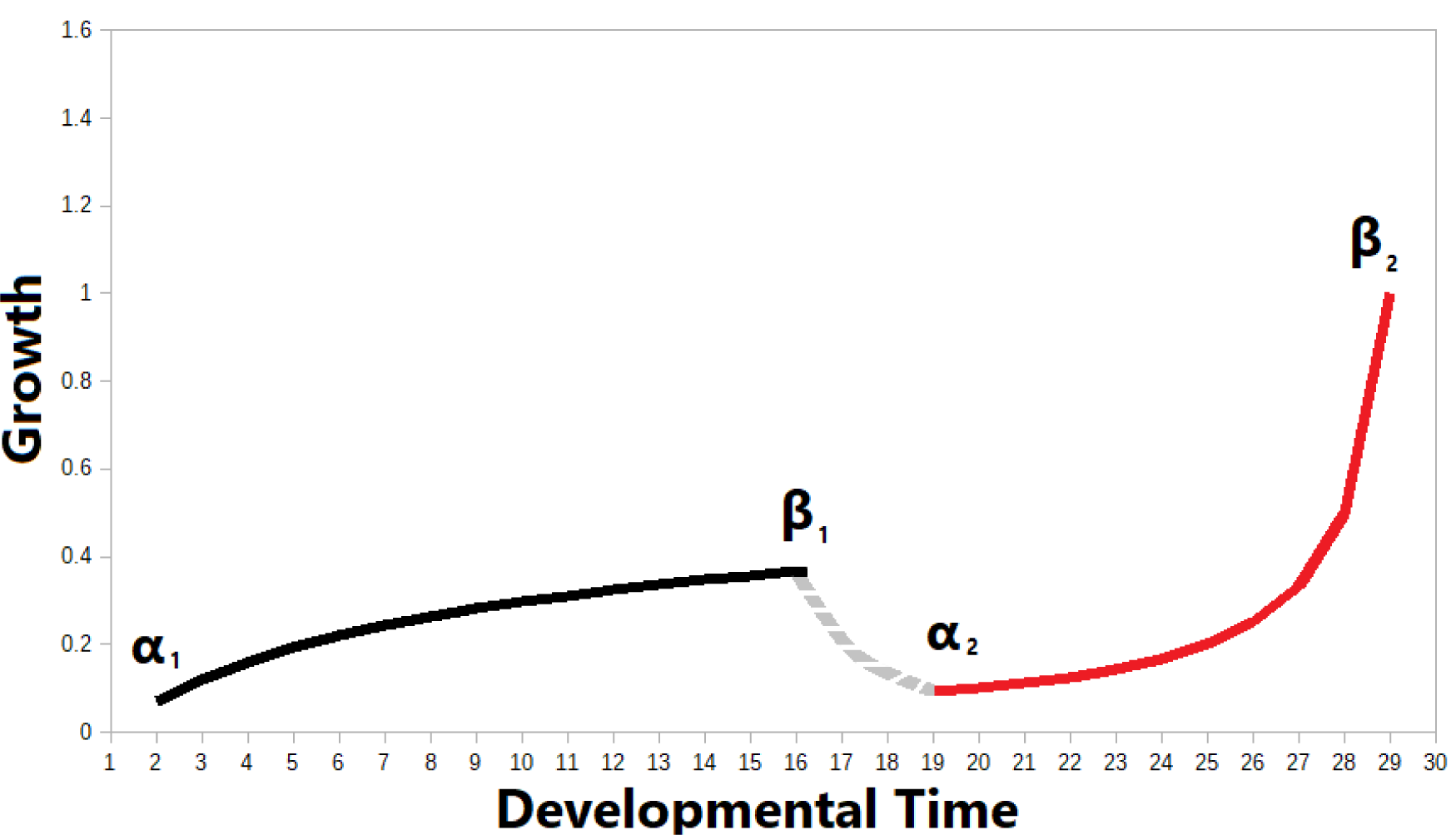
An example of a nonlinear switching event during developmental growth between two growth trajectories. Parameter α represents the start of the growth trajectory, and parameter β represents the end of this growth process. In this example of compound heterochrony, two heterochronic regimes are separated by a dedifferentiation and decellularization process. Gray dashed line between β_1_ and α_2_.

### Computational Regulatory Model

Binary GRNs model the expression of phenotypic change by modeling the network of heterochrony genes that produce an output of controlled growth. Each node in the tree represents a gene or regulatory element, and the tree structure itself is structured according to their relative causal effect on each other. With the addition of TSMs, epistasis can also be represented by introducing lateral branches across nodes at the same level. Structurally, the binary GRN model demonstrates the contingent nature of single factors and their effect on the collective regulation of growth. Each tree structure represents a single set of heterochrony genes for a specific developmental pathway [38]. Therefore, tree structures can be overlapped to simulate cross-talk between different networks of heterochrony genes.

Each node represents a hypothetical gene or regulatory element. Each level consists of a transformation that produces an output at the branch tips of the tree. As a result, children genes at a single level of the binary tree are considered to be a single functional unit (*cis*) of DNA. By contrast, when a gene at one level turns on or off a gene at the next level, it is said to act in *trans*. Lateral activation (within the same level of the tree) occurring between children pairs activate each other in *cis*. These overlapping structures, which result in a TSM, not only allow us to simulate switching behaviors, the aforementioned cross-talk, and nonlinear behaviors [42], but also result in emergent computational mechanisms.

In an overlapping example, the GRN maps the three types of behavior embedded in network arcs to numeric states: a state of “0” equals “off”, a state of “1” equals “on in -*trans*”, and a state of “2” equals “on in -*cis*”. Table 1 shows all possible ordinal paths and their corresponding states for an order 3 binary tree. Tables 2 and 3 show activity in the same tree over 10 timepoints for -trans and -cis pathways, respectively. GRNs can also be organized to show reciprocal connections between nodes. In such cases, the rules of the original tree hold, with the only difference being a potential regulatory mechanism between the parent and children. Figures 5 and 6 demonstrate these relationships, with Figure 5 demonstrating a GRN with reciprocal connections and Figure 6 providing an example of a reciprocally-connected GRN with a TSM.

**Table 1.** All pairwise ordinal paths (network arcs) in an order 3 binary tree. Ordinal path leads from source to destination.

| Source | Destination | State |
| --- | --- | --- |
| N/A | 0 | 1 |
| N/A | 0 | 1 |
| 0 | 00 | 1 |
| 0 | 01 | 1 |
| 1 | 10 | 1 |
| 1 | 11 | 1 |
| 01 | 10 | 1 |
| 0 | 1 | 2 |
| 00 | 01 | 2 |
| 10 | 11 | 2 |

**Table 2.**
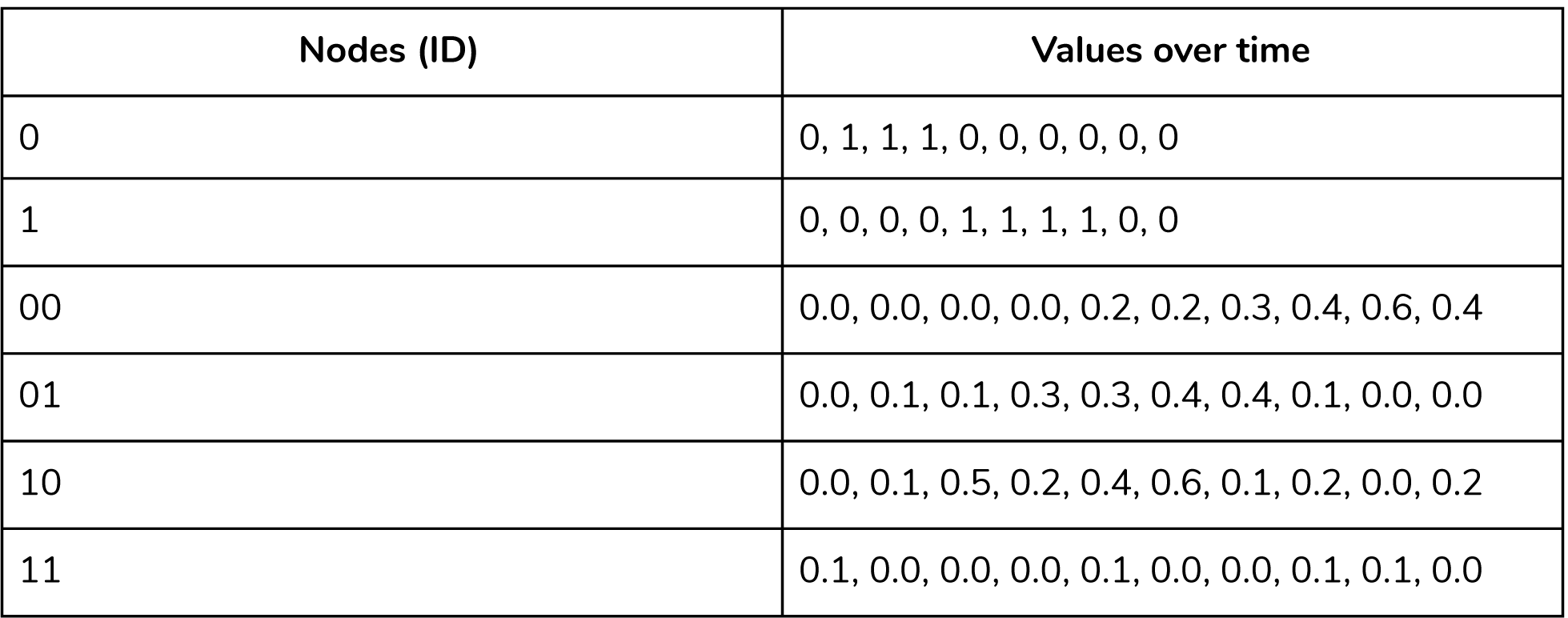
Origination nodes (on ordered tree) and values over time for activity in *trans*. Values for the master control gene (root of tree) are always 1. Examples are pseudo-data.

**Table 3.** Origination nodes and values over time for activity in *cis*. Examples are pseudo-data.

| Nodes (ID) | Values over time |
| --- | --- |
| 0 <--> 1 | 0, 1, 1, 0, 1, 0, 0, 0, 1, 0 |
| 00 <--> 01 | 0.0, 0.2, 0.3, 0.3, 0.5, 0.7, 0.4, 0.2, 0.1, 0.0 |
| 10 <--> 11 | 0.0, 0.0, 0.0, 0.0, 0.2, 0.2, 0.2, 0.2, 0.2, 0.0 |

**Figure 5.**
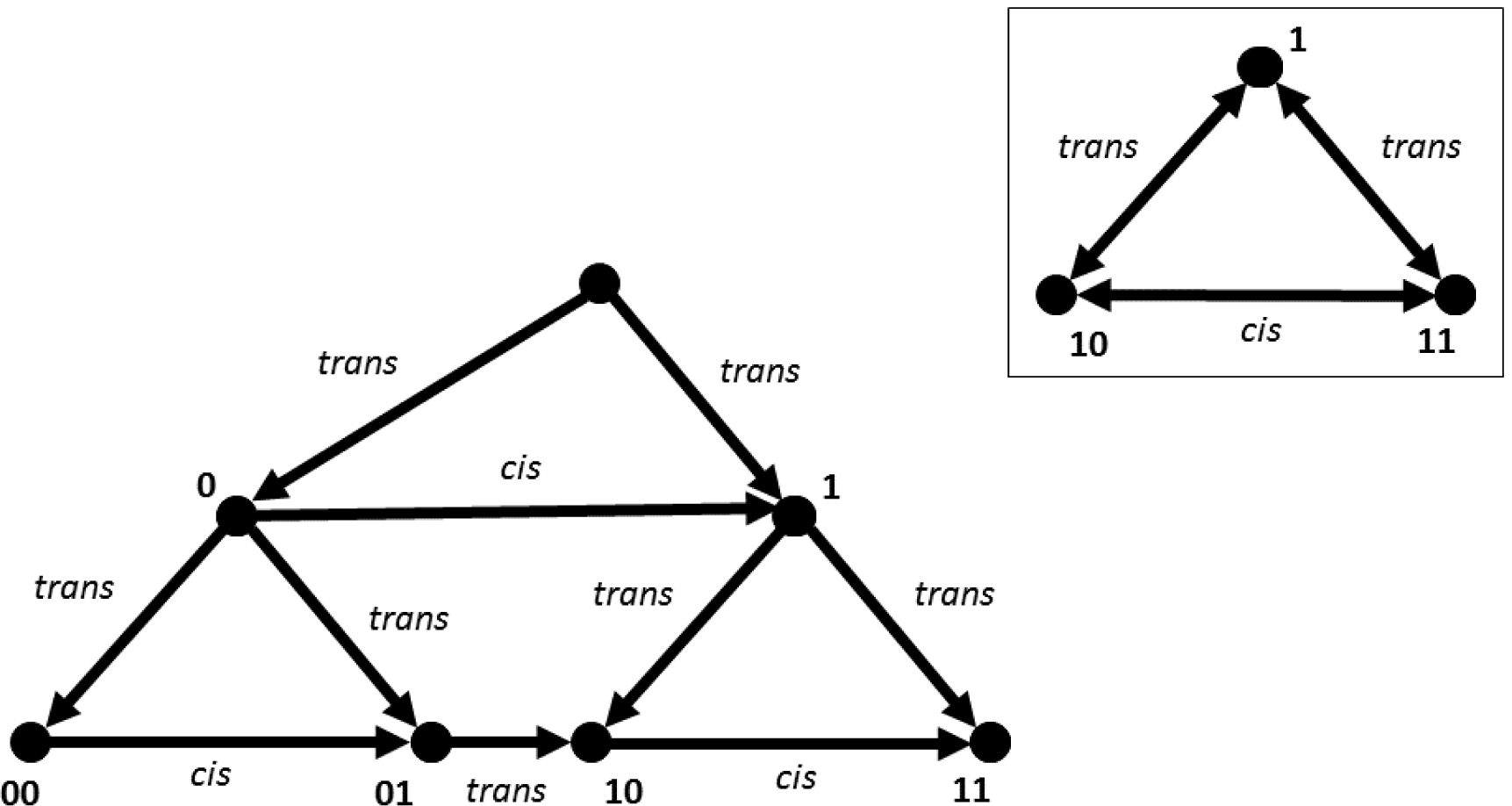
An order-3 GRN with nodes acting in both -cis and *trans*. INSET: an example of a TSM as network motif.

**Figure 6.**
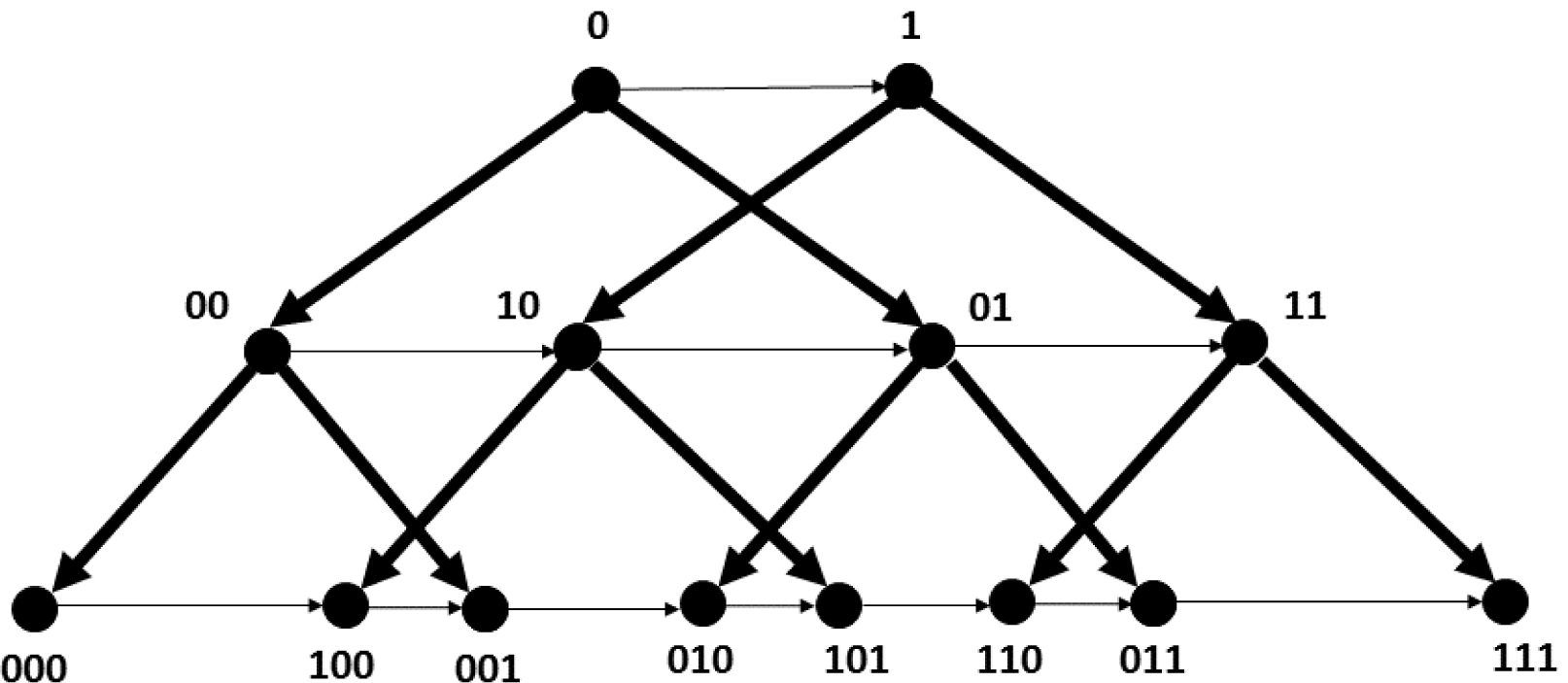
Two overlapping GRNs with overlapping nodes and ordered arcs forming a hierarchical compound model.

Each tree has an identity (0,1) that comes before the node identity (0,1). The leftmost node at order 3 in tree 0 is (000). An order 3 tree is classified as if it has four layers, but this order is discontinuous within a single tree depending on the amount of overlap between each tree. For an overlapping set of order 2 trees, nodes 00 and 01 belong to tree A and nodes 10 and 11 belong to tree B, but their order from left to right is 00, 10, 01, 11. This order is referred to as an overlapping order, and results in reversals, asymmetries, and spatially-restricted increases in complexity.

### Complex Phenotypes and TSMs

Due to the connectivity patterns of individual TSMs, particularly among overlapping GRNs, we expect to see complex phenotypes arise from these interactions. One such phenotypic pattern we expect to see are interference patterns similar to interference (Moire) patterns. This interference mechanism might reveal itself in the form of overlapping coloration patterns along the body, or the juxtaposition of differing connectivity patterns in a nervous system. The τ parameter can also play a role in determining what these interference patterns look like, whether they look more like a overlapping sets of concentric circles (negative values for τ) or the orderly meeting of two sets of lines at orthogonal orientations (e.g. a grid-like pattern, τ = 0).

### Galton board model of axial differential growth

To understand the mechanism behind GRN emissions a bit more, we can turn to a toy model called the Galton board [43,44]. A Galton board passes objects through a triangular array of pins, with the object ultimately coming to rest in a bin at the bottom of the board. The bins are arrayed in such a way that an unbiased board results in objects obeying the central limit theorem (accumulating in bins towards the center, while tending to avoid the bins along the edges of the board.

The behaviors demonstrated on a Galton board can also be used to simulate the spatially differentiated outputs of a GRN (Figure 7). To do this, we will utilize a generic binary tree that allows for nodal activation to be developmentally contingent without the complexity of overlapping topologies or mode of action. When the nodes in this binary GRN are uniform, the central limit theorem is observed. As with the Galton board, the emissions (in this case, components of the phenotype) tend to accrete in a symmetric fashion. There is mass around the center of the anterior-posterior (A-P) axis, and relatively little phenotypic accretion at the ends of the axis. If we make the binary GRN heterogeneous in terms of its internal topology, the emissions will be asymmetrical. The emissions will not only be spread out across the A-P axis, but also exhibit a non-normal statistical distribution.

**Figure 7.**
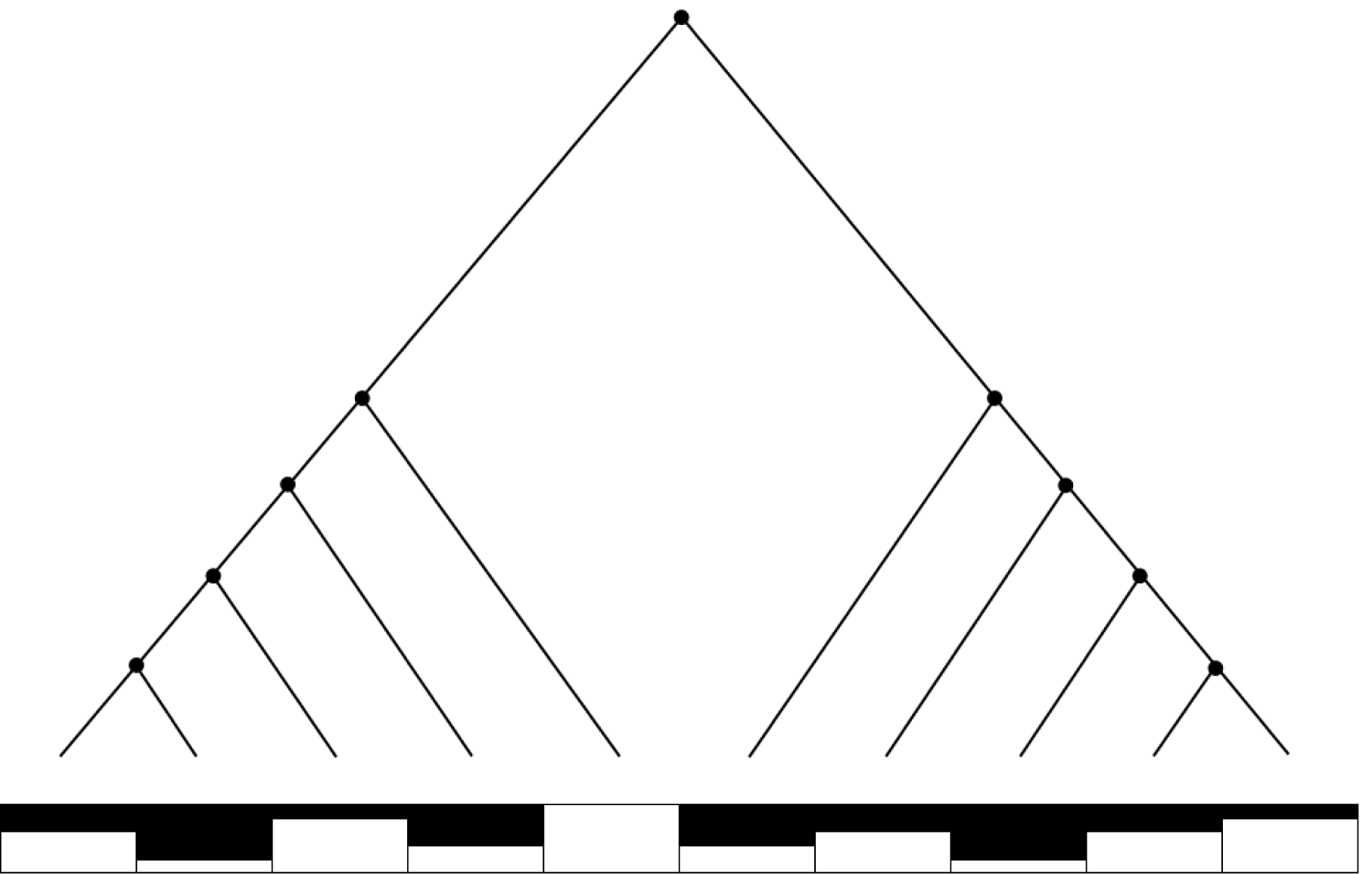
Binary GRN mapped to positions (bins) along the A-P axis. The Galton board is simulated by emitting a state from the root of the tree (top), which is activated or suppressed at each node and emitted at one of the tree’s terminal nodes (tips).

For the binary tree shown in Figure 7, each step in the accretion process follows a single path throughout the tree, and at the tips of the tree is emitted into a single bin in the series along the A-P axis. Outputs from the tree accumulate over time and result in a bimodal and asymmetric accretion distribution along the A-P phenotypic axis. Even though the topology is symmetrical, the nodes can be biased with a stochastic signal to result in various spatial patterns.

While the Galton board is always uniform with respect to time, differential behavior of the GRN can simulate the effects of temporal shifts in growth. Therefore, Figure 7 can also be understood in terms of parameters α, β, τ. Individual nodes can be turned on (α) and off (β) differentially, which can favor some internal paths over others. The onset of activation (α) can be delayed by τ, while the completion of growth/onset of quiescence (β) can be delayed by − τ. This leads to controlled differential accretion across the A-P axis.

### Mapping trees to radial differential growth

Figure 8 shows how the developing phenotype is characterized in a 2-D circular space. A differentiation tree [45] is constructed by sorting each tissue at each level and the relative expansion and contraction of each tissue spheres For a view of the animated process, see Supplemental Video 2. From the toy model demonstrated in this video, we can see a quantitative view of tissue differentiation. We can further understand our GRNs as something that produces a heterogeneous system where tissues are expressed in a time-dependent manner. While there are four parameters introduced in Supplemental Video 2 (*d*, θ, *c, v*^*g*^), we will pay attention only to *d* in connecting this toy model to our mathematical formulation of heterochrony.

**Figure 8.**
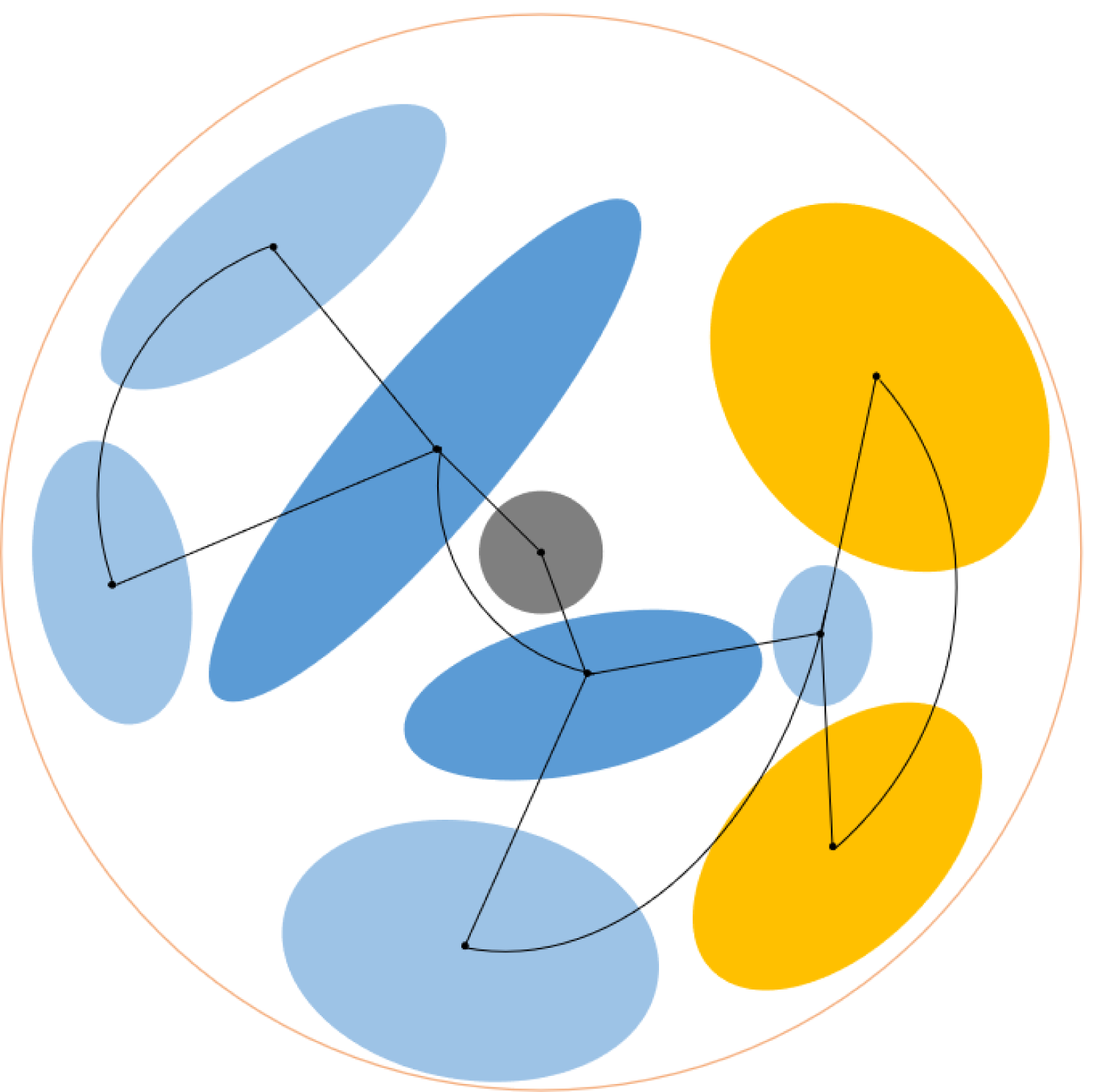
Expression tree with a hierarchical series of TSMs using a 2-D layered embryo. The tree extends radially from the center of the main sphere (embryo), and bifurcations in the tree result in a tissue. Layer colors, in order: gray (0), dark blue (1), light blue (2), yellow (3).

As with the Galton board example, we can map parameters α, β, and τ onto the radial differentiation tree. Figure 8 shows that the internal nodes of the tree serve as centroids for individual tissues, represented by a variety of circular and ovoid shapes. The radius of each tissue is determined by the onset and completion of growth with the delay serving to attenuate the uniformity of height and width. The onset of growth from a branch node (α) is always 0. The completion of growth (β) is some value *x* where *x*_*max*_ is the diameter of the embryo (*d*) divided by *2L* (number of levels). Attenuations in shape are determined by the vertical and horizontal components of τ : when τ _*vert*_ and τ _*horz*_ are both equal to 0, a circle results. When τ _*vert*_ is large and τ _*horz*_ is 0 (or vice versa), the more oblong a tissue becomes. From there, we can provide a fuller description of growth and form with respect to modes of GRN action (θ, *c*) and size scale (*v*^*g*^) of individual tissues as compared to the rest of the spherical tree.

## Discussion

In this paper, an approach involving a dynamic view of heterochrony has been advanced and a mechanism for the genomic expression of dynamic growth has been proposed. In terms of heterochrony, the basic mathematical structure of heterochrony theory has been extended to include multiple phases of growth. The genomic expression models introduced here utilize a representation of genetic regulatory mechanisms, particularly the cross-talk between pathways and modules, to produce outputs resulting in a growth trend. There are also several themes revealed in this paper. The first is the role of delays and differential time in the expression of phenotypes. Changes in phase across the developmental process are hypothesized to be due to multiple growth processes (expressed in the genotype) and changes in the rate and linkage of these growth processes [1-3,5,20]. A representation of the genotype also reveals how these dynamics might be regulated. This representation focuses on both regulation and gene-gene interactions [46] to produce a dynamical output. In cases where two binary GRNs overlap, it is suspected that TSMs provide a multitude of opportunities for local feedback. This may result in nonlinear outputs and other examples of complex regulation.

One overarching theme of this paper is that patterns of developmental growth and the progression of developmental dynamics are much more complex than assumed by contemporary theory. Using a combination of genotypic network representations and heterochronic scaling in the phenotype, we can begin to move towards viewing development as a highly complex and nonlinear process. Yet this approach also provides concrete mechanisms for guided generativity. In the case of *Drosophila* eye morphogenesis, switch-like behavior results from positive feedback between genes in the regulatory network [47]. Nonlinear positive feedback in the form of interacting positive feedback loops, sets the stage for dynamic bistability [48], or the conditions that enable switching mechanism responsible for both multiphasic and compound heterochrony. Dynamic bistability can be demonstrated in small and complex GRNs alike [49,50]. More generally, epigenetic landscapes [51] can be used to demonstrate switching as a function of differentiation and historical contingency. Future work will involve mapping simulations of development derived from our approach to developmental structures such as lineage trees and epigenetic landscapes.

## Acknowledgements

I would like to thank Aidan Rocke, Dr. Richard Gordon, members of Dynamics Days community, and members of the DevoWorm and Saturday Morning NeuroSim research groups for their feedback and discussion.

## Notes

https://figshare.com/articles/media/Supplemental_Files_for_Understanding_Developmental_Incongruity_Through_Dynamical_Heterochrony_/12743582

### Competing Interest Statement

The authors have declared no competing interest.

## References

[1] Alberch, P., Gould, S.J., Oster, G.F., and Wake, D. (1979). Size and shape in ontogeny and phylogeny. Paleobiology, 5(3), 296.

[2] McKinney, M.L. and McNamara, J.K. (2013). Heterochrony and the evolution of ontogeny. Springer.

[3] Smith, K.K. (2001). Heterochrony revisited: the evolution of developmental sequences. Biological Journal of the Linnean Society, 73, 169–186. doi:10.1006/bij1.2001.0535.

[4] Antonelli, P.L., Bradbury, R., Krivan, V., Shimada, H. (1993). A dynamical theory of heterochrony: Time-sequencing changes in ecology, development, and evolution. Journal of Biological Systems, 1(4), 451–487. doi:10.1142/S0218339093000264.

[5] Williamson, D.I. (1992). Larvae and Evolution: towards a new zoology. Chapman and Hall, New York.

[6] Technau, G. and Heisenberg, M. (1982). Neural reorganization during metamorphosis of the corpora pedunculata in *Drosophila melanogaster*. Nature, 295(5848), 405–407.

[7] Park, J.H. and Lee, G.G. (2018). Metamorphosis of the Central and Peripheral Nervous System (CNS and PNS) in Insects. Annals of Cell and Developmental Biology, 1(1), 1002.

[8] White, K., Pereira, A., and Cannon, L.E. (1983). Modulation of a neural antigen during metamorphosis in *Drosophila melanogaster*. Developmental Biology, 98(1), 239–244.

[9] Yamanaka, N., Rewitz, K.F., and O’Connor, M.B. (2013). Ecdysone control of developmental transitions: lessons from *Drosophila* research. Annual Review of Entomology, 58, 497–516.

[10] Consolas, C., Duch, C., Bayline, R., Levine, R.B. (2000). Behavioral transformations during metamorphosis: Remodeling of neural and motor systems. Brain Research Bulletin, 53(5), 571–583.

[11] Arlotta, P. and Berninger, B. (2014). Brains in metamorphosis: reprogramming cell identity within the central nervous system. Current Opinion in Neurobiology, 127, 208–214.

[12] Shepherd, D. and Bate, C.M. (1990). Spatial and temporal patterns of neurogenesis in the embryo of the locust (*Schistocerca gregaria*). Development, 108, 83–96.

[13] Ito, K. and Hotta, Y. (1992). Proliferation pattern of postembryonic neuroblasts in the brain of *Drosophila melanogaster*. Developmental Biology, 149, 134–148.

[14] Tissot, M. and Stocker, R.F. (2000). Metamorphosis in *Drosophila* and other insects: the fate of neurons throughout the stages. Progress in Neurobiology, 62, 89–111.

[15] Levine, R.B. (1984). Changes in Neuronal Circuits During Insect Metamorphosis. Journal of Experimental Biology, 112, 27–44.

[16] Huxley, J. (1932). Problems of Relative Growth. The Dial Press, New York.

[17] Christian, J.L. (2012). Morphogen gradients in Development: from form to function. Wiley Interdisciplinary Reviews: Developmental Biology, 1(1), 3–15.

[18] Watanabe, K., Wakita, J. Itoh, H., Shimada, H., Kurosu, S., Ikeda, T., Yamazaki, Y. Matsuyama, T., and Matsushita, M. (2002). Dynamical properties of transient spatio-temporal patterns in bacterial colony of *Proteus mirabilis*. Journal of the Physical Society of Japan, 71(2), 650–656.

[19] Zagorska-Marek, B. and Hejnowicz, Z. (1980). Discontinuous lines on the radial face of wavy-grained xylem as a manifestation of morphogenic waves in the cambium. Acta Societatis Botanicorum Poloniae, 49(1-2), 49–62.

[20] Williamson, D.I. (2003). The Origins of Larvae. Kluwer Academic Publishers, London.

[21] Costedoat, C., Pech, N., Chappaz, R., and Gilles, A. (2007). Novelties in Hybrid Zones: crossroads between population genomic and ecological approaches. PLoS One, 2(4), e357.

[22] Begun, D.J. et al. (2007). Population genomics: whole-genome analysis of polymorphism and divergence in *Drosophila simulans*. PLoS Biology, 5(11), e310.

[23] Liu, J., Yu, L., Arnold, M.L., Wu, C-H., Wu, S-F., Lu, X., and Zhang, Y-P. (2011). Reticulate evolution: frequent introgressive hybridization among Chinese hares (genus *Lepus*) revealed by analyses of multiple mitochondrial and nuclear DNA loci. BMC Evolutionary Biology, 11, 223.

[24] Figueiro, H.V. et.al (2017). Genome-wide signatures of complex introgression and adaptive evolution in the big cats. Science Advances, 3(7), e1700299.

[25] Baack, E.J. and Rieseberg, L.H. (2007). A genomic view of introgression and hybrid speciation. Current Opinion in Genetics and Development, 17(6), 513–518.

[26] Combes, M-C., Hueber, Y., Dereeper, A., Rialle, S., Herrera, J-C., and Lashermes, P. (2015). Regulatory Divergence between Parental Alleles Determines Gene Expression Patterns in Hybrids. Genome Biology and Evolution, 7(4), 1110–1121.

[27] Boto, L. (2010). Horizontal gene transfer in evolution: facts and challenges. Proceedings of the Royal Society B, 277, 819–827.

[28] Huang, J. (2013). Horizontal gene transfer in eukaryotes: the weak-link model. Bioessays, 35(10), 868–875.

[29] Soucy, S.M., Huang, J., and Gogarten, J.P. (2015). Horizontal gene transfer: building the web of life. Nature Reviews Genetics, 16, 472–482.

[30] Krotov, D., Dubuis, J.O., Gregor, T., and Bialek, W. (2014). Morphogenesis at criticality. PNAS, 111(10), 3683–3688.

[31] Mora, T. and Bialek, W. (2011). Are Biological Systems Poised at Criticality? Journal of Statistical Physics, 144, 268–302.

[32] Dubuis, J.O., Tkacik, G., Wieschaus, E.F., Gregor, T., Bialek, W. (2013). Positional information, in bits. PNAS, 110(41), 16301–16308.

[33] Verd, B., Crombach, A., and Jaeger, J. (2017). Dynamic maternal gradients control timing and shift-rates for *Drosophila* gap gene expression. PLoS Computational Biology, 13(2), e1005285.

[34] Verd, B., Clark, E., Wotton, K.R., Janssens, H., Jimenez-Guri, E., Crombach, A., and Jaeger, J. (2018). A damped oscillator imposes temporal order on posterior gap gene expression in *Drosophila*. PLoS Biology, 16(2), e2003174.

[35] Richard, J-P. (2003). Time-delay systems: an overview of some recent advances and open problems. Automatica, 39, 1667–1694.

[36] Artyomov, M.N., Meissner, A., and Chakraborty, A.K. (2010). A model for genetic and epigenetic regulatory networks identifies rare pathways for transcription factor induced pluripotency. PLoS Computational Biology, 6(5), e1000785.

[37] Moss, E.G. (2007). Heterochronic genes and the nature of developmental time. Current Biology, 17(11), R425–R434.

[38] Gordon, N.K. and Gordon, R. (2016). Embryogenesis Explained. World Scientific, Singapore.

[39] Fusco, G., Hong, P.S., and Hughes, N.C. (2014). Positional specification in the segmental growth pattern of an early arthropod. Proceedings of the Royal Society B, 281(1781), 20133037.

[40] Rios, L., Teran, J.M., Varea, C., and Bogin, B. (2019). Plasticity in the growth of body segments in relation to height-for-age and maternal education in Guatemala. American Journal of Human Biology, doi:10.1002/ajhb.23376.

[41] Klingenberg, C.P. (1998). Heterochrony and allometry: the analysis of evolutionary change in ontogeny. Biological Reviews Cambridge Philosophical Society, 73(1), 79–123.

[42] Brookings, T., Carlson, J.M., and Doyle, J. (2005). Three mechanisms for power laws on the Cayley tree. Physical Review E, 72(5), 056120.

[43] Limpert, E., Stahel, W.A., and Abbt, M. (2001). Log-normal Distributions across the Sciences: keys and clues. BioScience, 51(5), 341–352.

[44] Brehmer, J., Louppe, G., Pavez, J., and Cranmer, K. (2020). Mining gold from implicit models to improve likelihood-free inference. PNAS, 117(10), 5242–5249.

[45] Gordon, R. (1999). Hierarchical Genome And Differentiation Waves: novel unification of development, genetics and evolution. World Scientific, Singapore.

[46] Mensch, J., Carreira, V., Lavagnino, N., Goenaga, J., Folguera, G., Hasson, E., and Fanara, J.J. (2010). Stage-specific effects of candidate heterochronic genes on variation in developmental time along an altitudinal cline of *Drosophila melanogaster*. PLoS One, 5(6), e11229.

[47] Graham, T.G.W., Ali Tabei, S.M., Dinner, A.R., and Rebay, I. (2010). Modeling bistable cell-fate choices in the *Drosophila* eye: qualitative and quantitative perspectives. Development, 137, 2265–2278.

[48] Chang, D-E., Leung, S., Atkinson, M.R., Reifler, A., Forger, D., and Ninfa, A.J. (2010). Building biological memory by linking positive feedback loops. PNAS, 107(1), 175–180.

[49] Huang, S., Eichler, G., Bar-Yam, Y. and Ingber, D. E. (2005). Cell fates as high-dimensional attractor states of a complex gene regulatory network. Physical Review Letters, 94, 128701.

[50] Siegal-Gaskins, D., Grotewold, E. and Smith, G. D. (2009). The capacity for multistability in small gene regulatory networks. BMC Systems Biology, 3, 96.

[51] Ferrell, J.E. (2012). Bistability, Bifurcations, and Waddington’s Epigenetic Landscape. Current Biology, 22, R458–R466.

