## Supplemental File 1 for "Developmental Incongruity as a Dynamical Representation of Heterochrony"

### Modeling Heterochrony with Delays and Critical Periods

**Bradly Alicea**

**Orthogonal Research and Education Lab, OpenWorm Foundation**

### Growth Model of Heterochrony

Review of the Alberch et.al heterochrony model (1979). Their simple growth law can be stated as

$$\frac{dy}{da} = \begin{cases} 0, & \mathrm{a} < \alpha \\ ky, & \alpha < \mathrm{a} < \beta \\ 0, & \mathrm{a} > \beta \end{cases} \tag{1}$$

where *y* grows according to

$$y(a) = y_0 e^k a \tag{2}$$

between onset age $\alpha$ and offset age $\beta$.

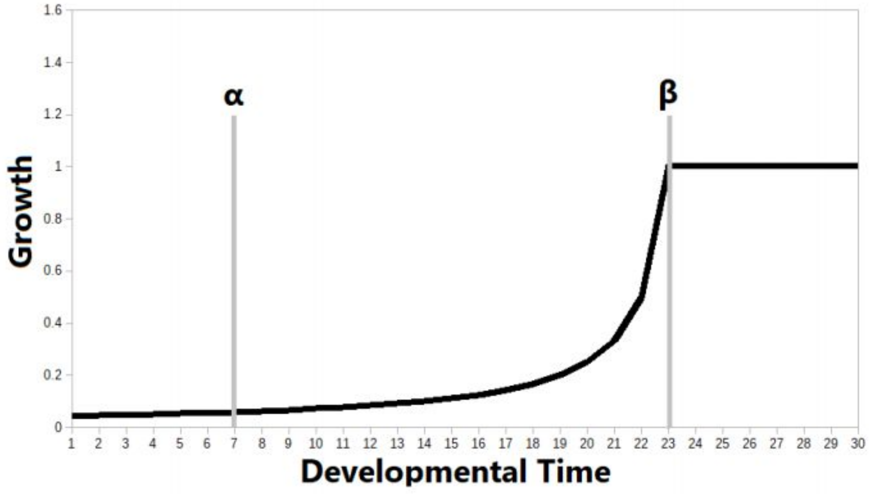


In this above example, growth is initiated at $\alpha$ and terminates at $\beta$, resulting in maintenance of an adult size.

The allometric growth law is a variant of the simple growth law, and involves two size/shape programs in the same orgaism.

For growth rates $k_1, k_2$, both $y_1$ and $y_2$ grow proportionally according to

$$y_1(t) = \lambda y_2(t)^b \tag{3}$$

where $b = \frac{k_1}{k_1}, \lambda = \frac{y_1(0)}{y_2(0)}$ and $g$ is transformation (or change in growth rate). In general, $\frac{k_1}{k_0} \sim g_1$

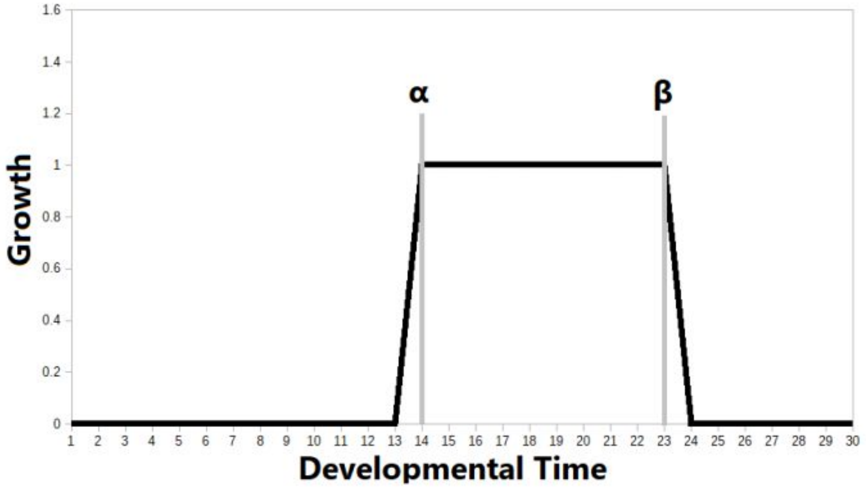

In the example above, growth is initiated at $\alpha$ and terminates at $\beta$, resulting in finite growth.

In the first example below, the red trajectory represents neoteny, or slower growth relative to baseline (where $k$ represents the slope, or rate of growth).

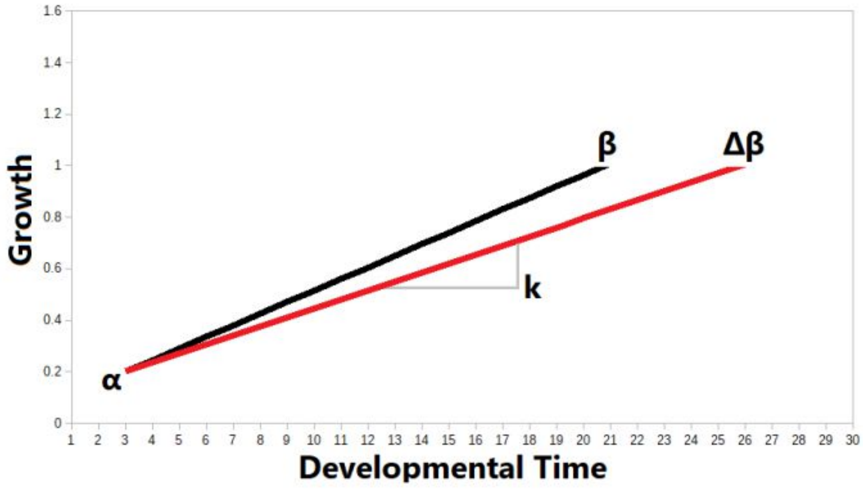


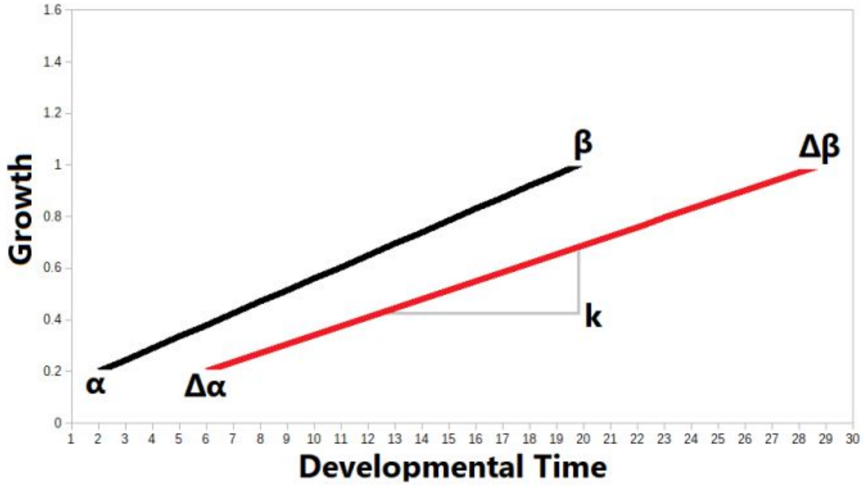


In the example above, the red trajectory also represents neoteny, but is caused by both slower growth and delayed onset of growth relative to baseline (where $k$ represents the slope, or rate of growth). McKinney and McNamara (2013) have developed a typology of terms for shifts in growth trajectory and rate due to heterochrony.

### Sequence Heterochrony

An alternate version of heterochrony comes from Smith (2001) and is called "sequence heterochrony". In sequence heterochrony, the developmental trajectory is treated as a series of developmental events. These events shift in time relative to one another so that they can overlap or occur separate depending on the species.

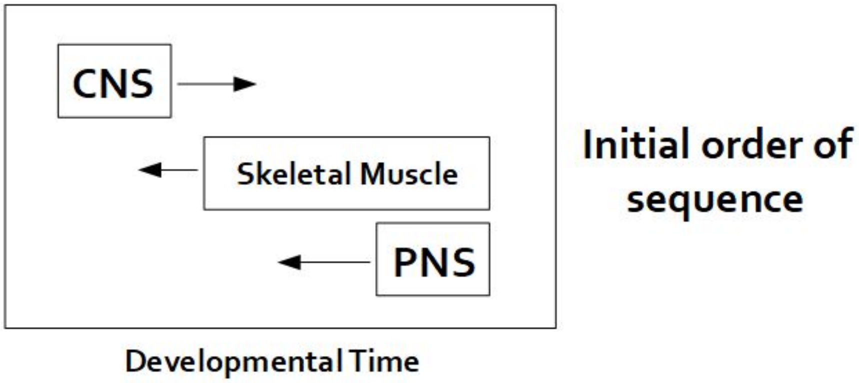


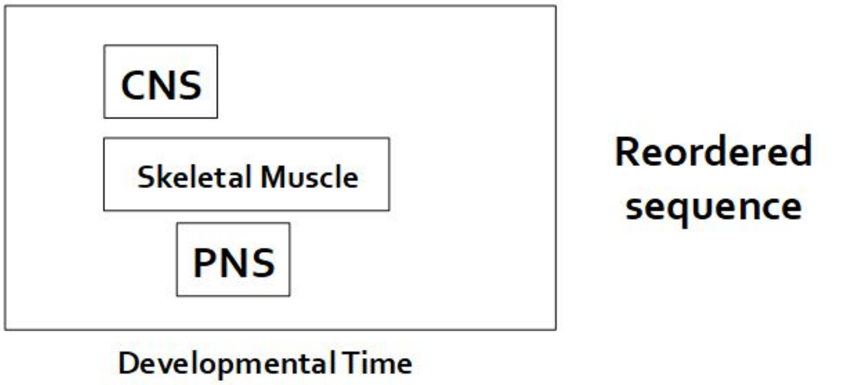


### Delay Model of Heterochrony

Delay Differential Equations (DDEs) are characterized by time-delay systems in Richard (2003). In their general form, a time-delay system is

$$\frac{d}{dt}x(t) = f(t, x(t), x_t) \qquad (4)$$

where $x_t = [x(\tau) : \tau \leq t]$ represents the trajectory of a solution in the past.

Both DDEs and time-delay systems can be solved using the method of steps (dde23 solver in Matlab). For a DDE with a single delay, the equation can be structured as

$$\frac{d}{dt}x(t) = f[x(t), x(t-\tau)] \tag{5}$$

We can also characterize time delays more specifically with respect to heterochrony and developmental growth trajectories. In equation (1), $\alpha$ and $\beta$ both exhibit systematic time delays ($\tau$): $\alpha(\tau)$, $\beta(\tau)$.

Delay in the growth trajectory is characterized over the interval ( $\alpha$, $\beta$), and is equivalent to $\Delta k$. The total length of the delayed process is $(\beta + \tau) - (\alpha + \tau)$. Using this formulation, the rate of a delay process is $k_0$, the delay rate is $\frac{k_1}{k_0}$, and the length of a delay process is $\beta_x - \alpha_x$

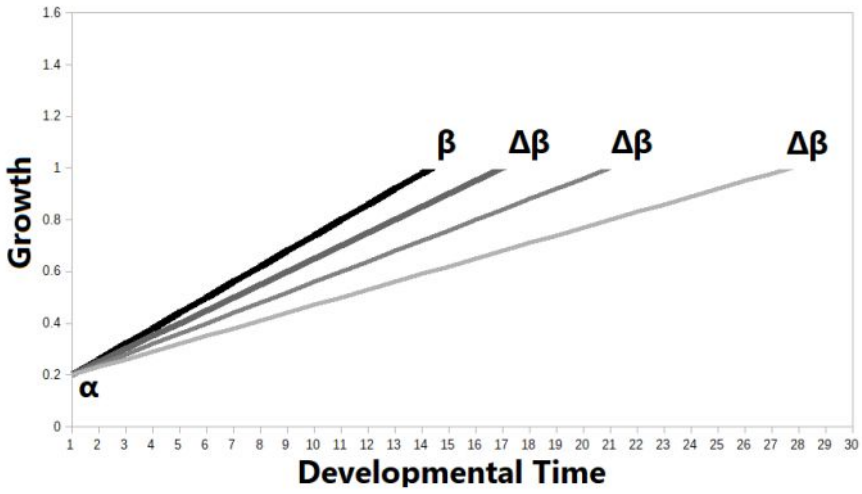


### Compound Heterochrony

Now that we have discussed heterochrony, allometry, and growth, we can discuss *compound heterochrony*. Recall from equation () that allometry is a measure of relational growth between two organs or phenotypic modules. Compound heterochrony involves contributions to growth exhibited between two developmental programs expressed in the same organism (see Williamson (1992). Unlike in the allometry example, compound heterochrony is usually not relational. Rather, the component programs are either cooperative or competitive in their effects on the organism's developmental trajectory.

We will consider two specific cases of compound heterochrony:

A) two developmental programs expressed in either a sequential or overlapping fashion with respect to time.

B) two developmental programs expressed in an overlapping fashion in a way that results in two sets of developmental parameters for growth ($y_1$ , $y_2$).

In the example of compound heterochrony in the figures below, the blue rectangle is the "centipede" program, while the red triangle is the "butterfly" program.

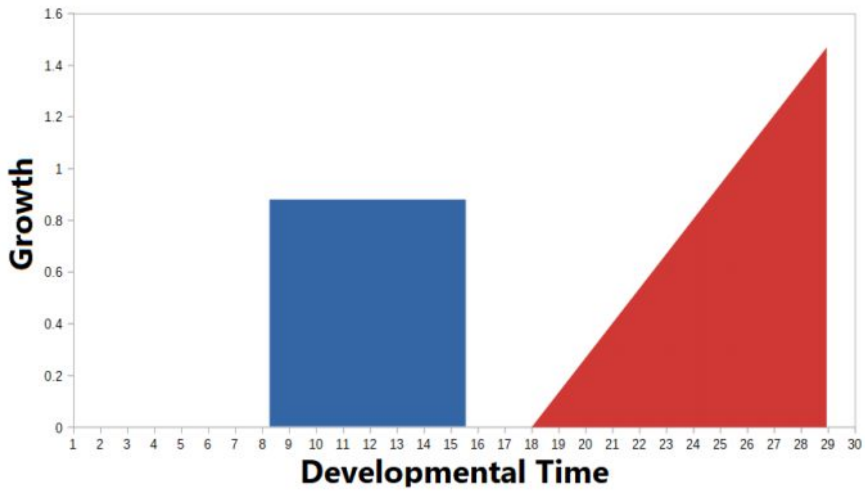

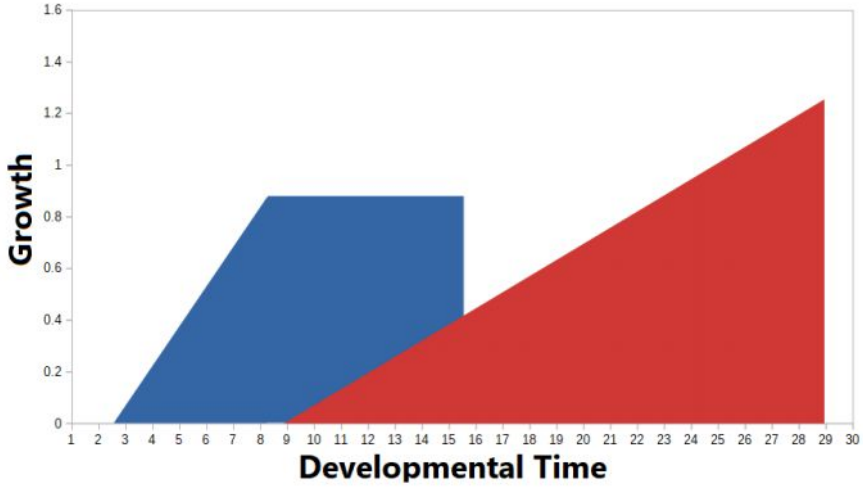


In the example of compound heterochrony in the figure above, Program 1 is blue and Program 2 is red. In compound heterochrony, $y_1$ and $y_2$ are nominally independent ($y_1 \perp y_2$).

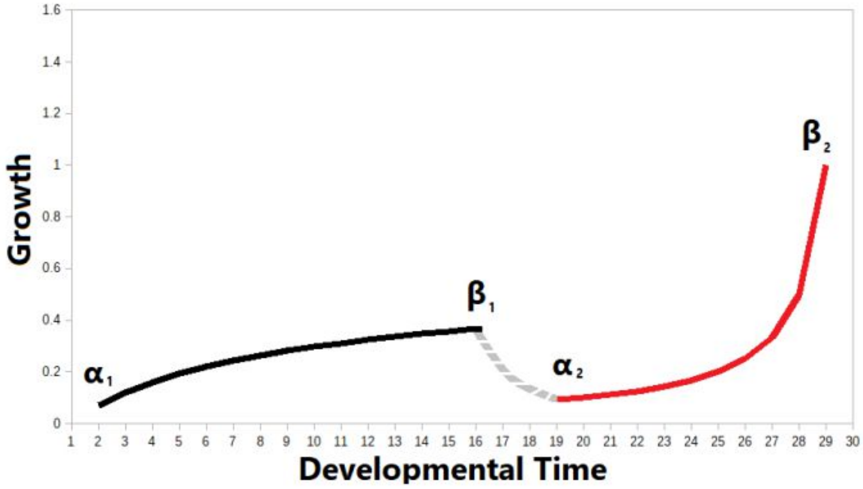


In the example of compound heterochrony in the figure above, two developmental programs are separated by a dedifferentiation and decellularization process (gray dashed line between $\beta_1$ and $\alpha_2$).

In the example of compound heterochrony in the figure below, overlapping developmental programs $\alpha, \beta$ (black) and $\Delta\alpha, \Delta\beta$ (red). The overlap of developmental programs are defined by a delay in onset ($\tau_\alpha$) and a delay in termination ($\tau_\beta$)

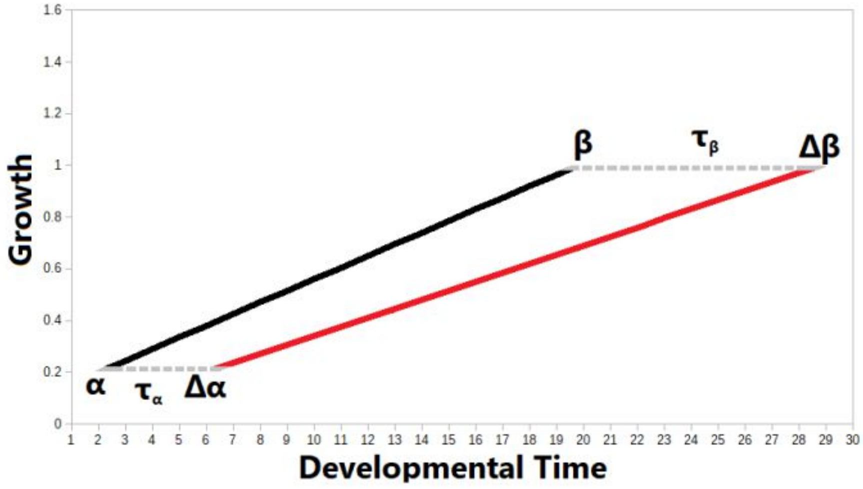


Generally speaking, the delay in program onset can be described as

$$\alpha_2 - \alpha_1 = \tau_\alpha \tag{6}$$

$$\tau_\alpha = \begin{cases} if \alpha_2 > \beta_1, & \text{transformation} \\ if \alpha_2 < \beta_1, & \text{superimposition} \end{cases} \tag{7}$$

while the delay in program termination can be described as

$$\beta_2 - \beta_1 = \tau_\beta \tag{8}$$

$$\tau_\beta = \begin{cases} if k_2 > k_1, & \text{programs more integrated} \\ if k_2 < k_1, & \text{programs less integrated} \\ if k_2 = k_1, & \text{programs complementary} \end{cases} \tag{9}$$

Overall, delayed growth is a function of three parameters

$$x(g) = f(\tau, \Delta t, \theta) \tag{10}$$

where $\tau$ is the delay in *t*, $\Delta k$ is change in slope, and $\theta$ is the phase angle between $f(g)$ and $f(g)'$, or between $f(g)$ and a baseline.

This can be restated as an evolution equation

$$u' = f(t, u), u'' = f(t, u, u) \tag{11}$$

where $u(t)$ is a growth function. This can be used in cases where an exact solution to the developmental trajectory is desired.

We may also observe switching from one developmental program to another. In the figure below, the two developmental growth trajectories are not only nonlinear, but also intersect at the termination point for one of the trajectories (black). This violates the cases for $\tau_\beta$ described in equations (6-9), but also provides a bifurcation point thyat enables a set of recombinant trajectories.

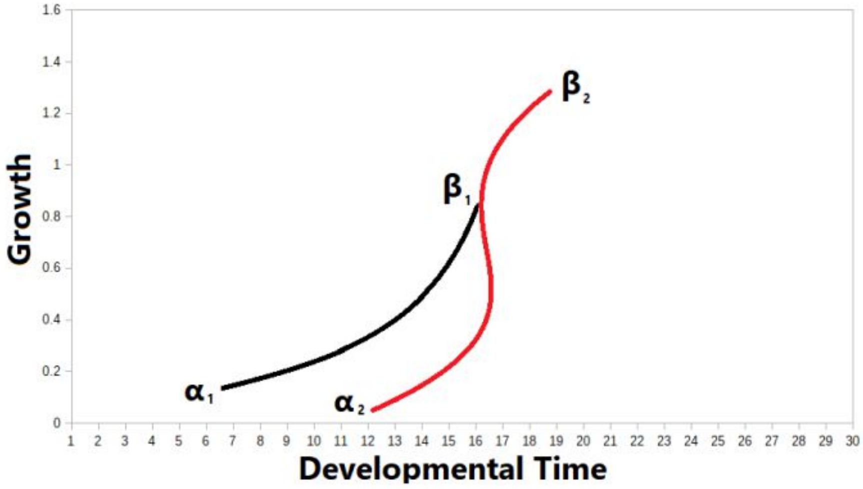


### Heterochrony and the Critical Period

The simple growth model described in equations (1-3) is not explicitly constrained by a temporal window. Life history theory (Promislow and Harvey, 1990) posits that the juvenile stage is a finite period, subject to modification. We can characterize these limits to growth by applying Hensch's (2005) model of critical periods in development. Multiple critical periods for multiple modules and/or functions.

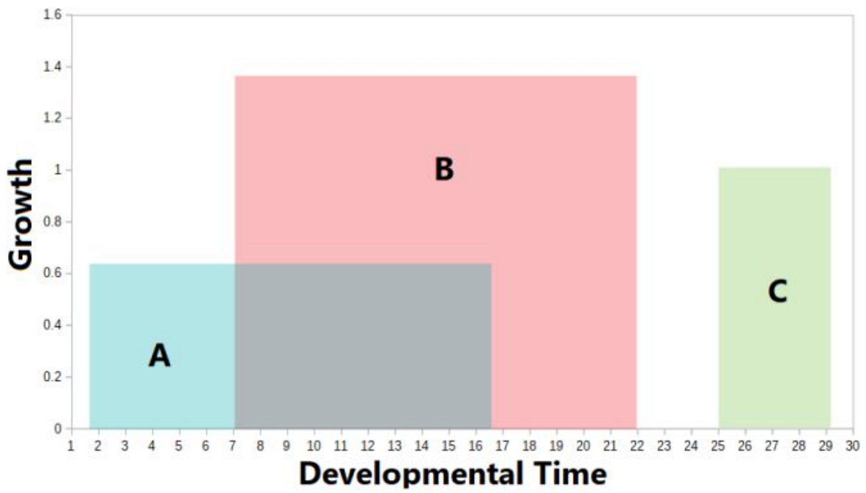


In general, we can describe a critical period window as

$$\omega(g,t) = min(g,t) \rightarrow max(g,t) \tag{12}$$

where $g$ is growth and $t$ is developmental time. We can also relate the simple growth to the critical period model in the following manner:

a) critical period window is $\Delta t$ wide ($t_1 - t_0$) and $p$ tall (where $p = p_0 \rightarrow \max_p$). In general, $g(x) \propto p(x)$.

b) growth decelerates ($\beta$ gets moved back in time).

c) new position of $\beta$ with respect to time ($\beta$') exceeds the length of the critical period window (at $t_1$).

d) this interaction prematurely terminates growth. The difference in growth between $\beta$ and $\beta$' is ($\Delta g$).

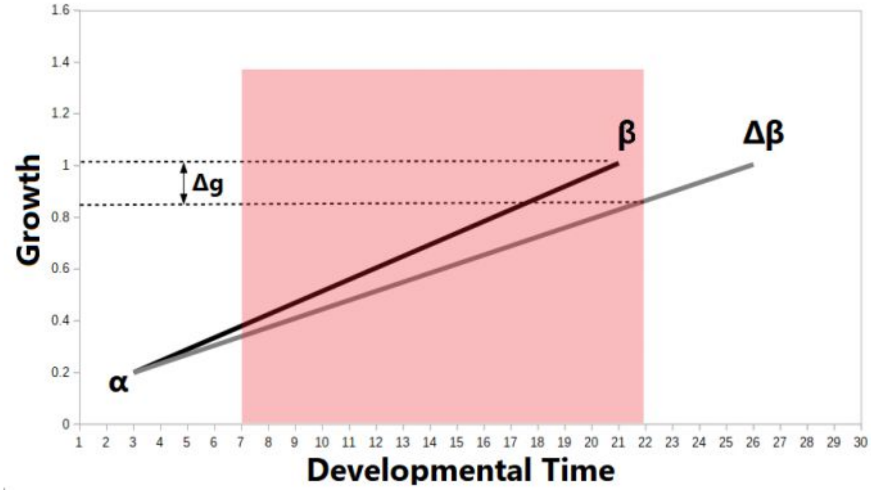


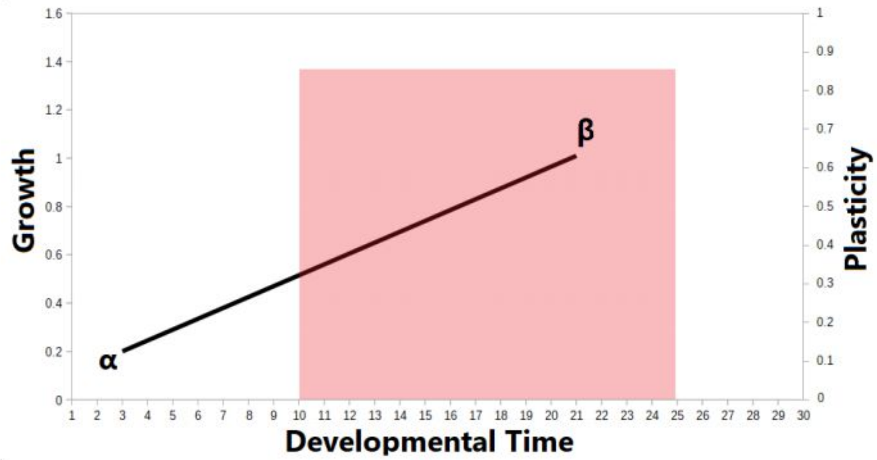


### Heterochrony and Dynamical Systems

One way to characterize growth trajectories using a dynamical approach is to approximate the rate of separation between an initial condition and evolution of system $Z$ at time $t$. In this case, $\alpha$ serves as the initial condition, while $\beta$ and $\Delta\beta$ serve as alternate trajectories with termination at time $t$. This distance is measured between two trajectories, which can either be linear or nonlinear.

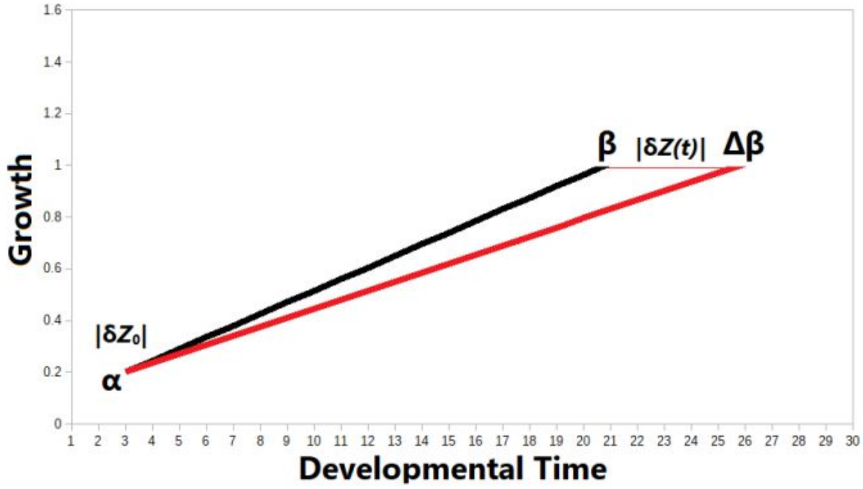


In the above example, $\Delta\beta$ is determined by a delay. In the example below, $\Delta\beta$ is determined by both a partial delay (delay in the earlier developmental program but not the later one) and the end of the critical period window. In both cases, the rate of divergence between both growth trajectories is measured by $|\delta Z(t)|$.

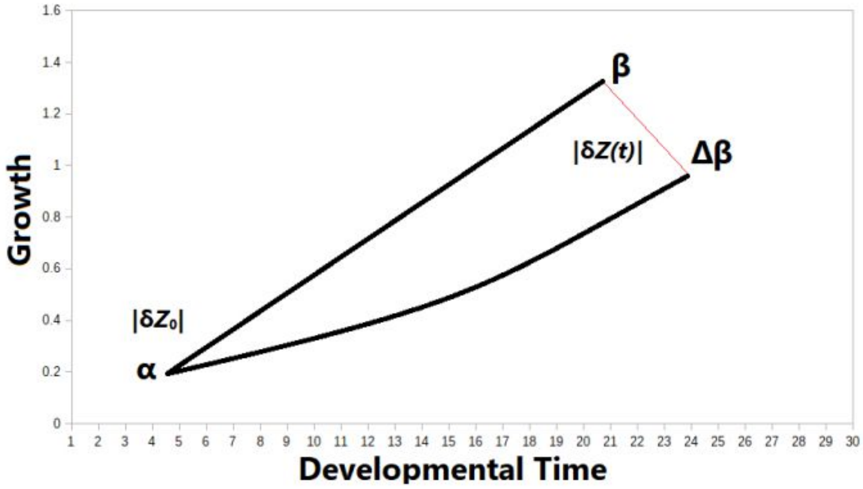


In cases of compound trajectory where there are two intersecting developmental programs, the initial condition is a bifurcation point $\alpha_2$ at some point in the growth trajectory rather than the initial condition $\alpha_1$. In this case, we observe at least two developmental programs producing alternate growth trajectories. These growth trajectories are also linked to a common history, or the same developmental program over interval $(\alpha_1, \alpha_2)$.

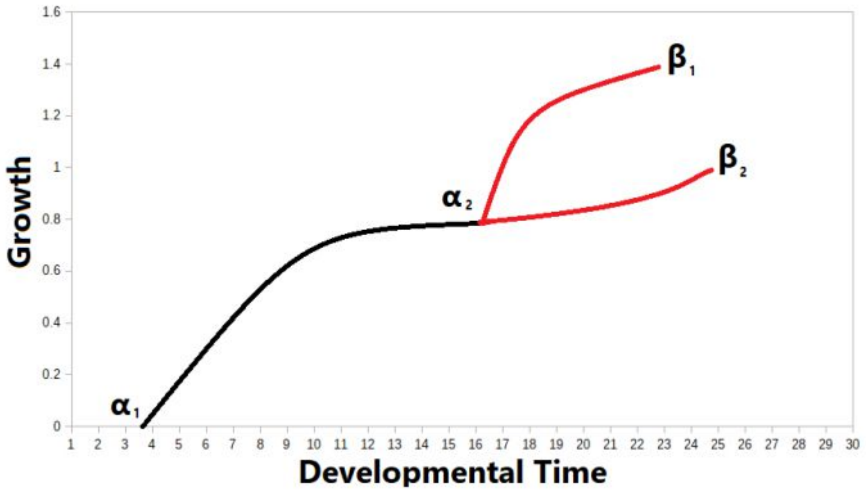


### References

Alberch, P., Gould, S.J., Oster, G.F., and Wake, D. (1979). Size and shape in ontogeny and phylogeny (https://www.researchgate.net/publication/272152220_Size_and_s *Paleobiology*, 5(3), 296.

Hensch, T.K. (2005). Critical period plasticity in local cortical circuits (https://www.nature.com/articles/nrn1787). *Nature Reviews Neuroscience*, 6(11), 877–888.

McKinney, M.L. and McNamara, J.K. (2013). Heterochrony and the evolution of ontogeny (https://books.google.com/books/about/Heterochrony.html?id=AWrIBwAAQBAJ&source=kp_book_description). Springer.

Promislow, D.E.L. and Harvey, P.H. (1990). Living fast and dying young: A comparative analysis of life-history variation among mammals

(https://zslpublications.onlinelibrary.wiley.com/doi/abs/10.1111/j.14
7998.1990.tb04316.x). *Journal of Zoology*, 220, 417-437.

Richard, J-P. (2003). Time-delay systems: an overview of some recent advances and open problems
(https://www.sciencedirect.com/journal/automatica/vol/39/issue/10
*Automatica*, 39, 1667-1694.

Smith, K.K. (2001). Heterochrony revisited: the evolution of developmental sequences
(https://onlinelibrary.wiley.com/doi/abs/10.1111/j.1095-8312.2001.tb01355.x). *Biological Journal of the Linnean Society*, 73, 169-186. doi:l0.1006/bij1.2001.0535.

Williamson, D. (1992). Larvae and Evolution: towards a new Zoology
(https://books.google.com/books/about/Larvae_and_Evolution.htm
id=MV8WAQAAIAAJ). Springer.
